# Does Inter-Protein Contact Prediction Benefit from Multi-Modal Data and Auxiliary Tasks?

**DOI:** 10.1101/2022.11.29.518454

**Authors:** Arghamitra Talukder, Rujie Yin, Yuanfei Sun, Yang Shen, Yuning You

**Affiliations:** Department of Electrical and Computer Engineering, Texas A&M University, USA

## Abstract

Approaches to *in silico* prediction of protein structures have been revolutionized by AlphaFold2, while those to *predict interfaces between proteins* are relatively underdeveloped, owing to the overly complicated yet relatively limited data of protein–protein complexes. In short, proteins are 1D sequences of amino acids folding into 3D structures, and interact to form assemblies to function. We believe that such intricate scenarios are better modeled with additional indicative information that reflects their multi-modality nature and multi-scale functionality. To improve binary prediction of inter-protein residue-residue contacts, we propose to augment input features with multi-modal representations and to synergize the objective with auxiliary predictive tasks. (**i**) We first progressively add three protein modalities into models: protein sequences, sequences with evolutionary information, and structure-aware intra-protein residue contact maps. We observe that **utilizing all data modalities delivers the best prediction precision**. Analysis reveals that evolutionary and structural information benefit predictions on the difficult and rigid protein complexes, respectively, assessed by the resemblance to native residue contacts in bound complex structures. (**ii**) We next introduce three auxiliary tasks via self-supervised pre-training (binary prediction of protein-protein interaction (PPI)) and multi-task learning (prediction of inter-protein residue–residue distances and angles). Although PPI prediction is reported to benefit from predicting inter-contacts (as causal interpretations), it is not found vice versa in our study. Similarly, the finer-grained distance and angle predictions did not appear to uniformly improve contact prediction either. This again reflects the high complexity of protein–protein complex data, for which **designing and incorporating synergistic auxiliary tasks remains challenging**.

## 1 Introduction

Proteins are essential biomolecules of life. The seemingly simple, linear bio-polymers made of 20 standard amino acids can adopt vast choices of 1-dimensional (1D) amino-acid sequences, fold into unique 3-dimensional (3D) tertiary structures, interact with each other to form protein complexes and assemblies, and perform various molecular functions underlying cellular processes.

Intrinsically multi-modality, protein data are increasingly treated that way in machine learning. Some of the most significant examples come from the field of protein structure prediction. CASP (Critical Assessment of Structure Prediction) 13 in 2018 proved the success of predicting 2-dimensional (2D) intra-protein residue–residue contact/distance maps as images from 1D sequences, including pioneering RaptorX-Contact [1], AlphaFold [2], and C-I-TASSER [3]. CASP 14 witnessed giant leaps by AlphaFold2 [4] that combines transformers for 1D sequences, message passing for 2D residue–residue contact maps, and roto-translationally equivariant neural networks for 3D atomic coordinates to jointly model multiple modalities of proteins in an end-to-end fashion. Inspired by AlphaFold2, RoseTTAFold [5] also followed the spirit of multi-modal end-to-end learning.

Unlike protein structure prediction, the sibling field of protein complex/assembly structure prediction has yet seen as successful technology breakthroughs, largely due to more complexity and less data. The straightforward extension from intra- to inter-protein contact prediction using Complex Contact [6] did not reach the full potential, partly because paired multiple sequence alignments as feature sources and paired structures as label sources are much less abundant than their unpaired counterparts. The straightforward extension from tertiary to quaternary structure prediction using RoseTTAFold or AlphaFold2, by artificially inserting amino acids between two proteins [5, 7], saw limited success. Recently, the inter-protein contact prediction field saw some exploration into the cross-modality, multi-task learning and pre-training. A state-of-the-art (SOTA) method [8] utilizes as input the coevolutionary features (generated from the MSA (multiple sequence alignment) transformer) and intra-protein distance maps (generated from predicted or experimental structure information). It then combines a deep residual network, a channel-wise attention mechanism, and a spatial-wise attention mechanism to predict the interchain distance maps of both homodimers and heterodimers. These studies naturally lead us to explore the possible synergy of more tasks (such as distance prediction) besides inter-protein residue contact prediction as well as the systematic studies of input information from more modalities besides protein sequences.

### Contributions

With the aforementioned review of the field, we are well-motivated to dive into the following two-fold question: *Can one improve inter-protein residue contact prediction via incorporating multi-modal data and synergizing multiple predictive tasks?*

Our first effort is to examine the effect of multi-modal protein data, coupled with the SOTA feature extractors. Motivated from [8, 9], three modalities are included: (i) protein sequence representations extracted from HRNN [10], (ii) sequences with evolutionary information encoded by an in-house protein language model, similar to ProtBERT [11], pretrained using protein domain sequences from Pfam RP15; and (iii) structure-aware intra-protein residue contact maps with GAT encoders [12]. Our experiments uncover that, **by adding extra modalities to models, we can boost the predictive performance** (AUPRC) from sequence-only to all modalities by 34.37%, surpassing the SOTA performance by 26.47% [13]. Moreover, we categorize test protein complexes into three cases based on the levels of the unbound-to-bound protein conformational changes [14], and observe that evolutionary and structural information benefit predictions on the rigid protein complexes by 41.4% and 69%, respectively; and they did on the flexible/difficult cases by 83.3% and 66.6%, respectively.

Next, we explore on how auxiliary predictive tasks could impact inter-protein residue contact prediction. Since it is reported that protein-protein interaction (PPI) prediction benefits from contact prediction acting as a casual interpretation [15, 9], we first assay whether it is vice versa (via pre-training). Moreover, we enforce models to simultaneously predict inter-protein residue-residue distances and angles, which are closely related to (actually finer details about) binary contacts and presumably aid the prediction. Unfortunately, **none of the auxiliary tasks are shown to provide synergistic performance gains**, on protein complexes across all categories.

## 2 Method

### Tasks

(1) *Target task*. We target at predicting the binary, inter-protein residue contact map (or inter-contact in short) given two proteins as inputs. Specifically, considering two proteins of certain representations containing *L*_1_ and *L*_2_ residues, respectively, a model is asked to output a interface matrix *C*_pred_ ∈ [0,1]^*L*_1_ × *L*_2_^ whose entries stand for the probabilities of the inter-contact between two corresponding residues across proteins. (2) *Auxiliary tasks*. Besides binary prediction of inter-contacts, we also have two types of auxiliary tasks: binary prediction of PPI, the consequences of inter-contacts (incorporated through pre-training) and multi-classification of inter-protein 2D geometries, the finer details of inter-contacts (incorporated through multi-task learning concurrent with binary contact prediction). Specific 2D geometries follow [5], including *C_β_*–*C_β_* distance, a dihedral angle along the virtual bond connecting two *C_β_* atoms, dihedral angles *θ_ij_* and *θ_ji_* defined along consecutive *N*–*C_α_–C_β_–C_β_* atoms, and planar angles *ϕ_ij_* and *ϕ_ji_* defined along consecutive *C_α_–C_β_–C_β_* atoms.

### Data

Considering the different mechanisms to incorporate the aforementioned two types of auxiliary tasks, there are two datasets.

#### Target task and auxiliary task of distance and angle prediction (multi-task learning)

We used the data from Docking Benchmark Version 5 (DB5) [16] and the split from [13]. Due to the extremely imbalanced ratio of positive and negative residue-contact pairs in interface contact maps (less than 1:1000), during training we down-sample negative contact pairs and leave the positive (native) ones intact [17] so that the positive-to-negative ratio reaches 1:10. Protein sequences are directly retrieved from PDB files and and per-residue featurization follows [13]. Intra-contact graphs are constructed from the unbound structures (PDB files) following the *k* nearest neighbor (*k*-NN) principle where *k* = 20. We derive the labels from the bound structures, including binary contacts and discretized distances and angles. More details are provided in Appendix A.

#### (2) Auxiliary task of PPI prediction (pre-training)

We curate 4243 positive interactions from [18] and 4856 negative interactions from [19]. To remove redundancy for PPIs, data are filtered into 1007 unique proteins for both positive and negative interactions. Protein sequences are retrieved from UniProt. We use random 60&/15%/25% split for training/validation/test sets.

### Encoders

We introduce three protein modalities encoded with SOTA neural networks. (**i**) Protein sequences are encoded through hierarchical recurrent neural networks (HRNN) [20, 21], (**ii**) sequences are additionally encoded using our own protein language model (pretrained using RP15 sequences) that captures evolutionary information, and (**iii**) structure-aware intra-contact maps are embedded with graph attention network (GAT) [12]. Three resulting embeddings are then concatenated as the protein representation. The detailed architecture is depicted in Appendix B.

### Model Architecture and Training

The overall pipeline of our experiments is presented in Figure 1. In brief, we feed the model with multi-modal protein representations, encoded by the state-of-the-art (SOTA) neural networks. Afterwards, we introduce auxiliary predictive tasks through via pre-training or multi-task learning. During training, we use the balanced binary cross entropy loss for optimization which is weighted summed with the *ℓ*_2_-norm regularization loss to avoid overfitting. All models are trained with batch size of 2 and Adam optimizer for up to 200 epochs. Three hyper-parameters are tuned based on validation AUPRC: learning rate from {0.00001, 0.0001, 0.001}), GAT layer number from {1, 2, 3, 4} and dropout rate from {0.1, 0.2, 0.3, 0.4}).

**Figure 1:**
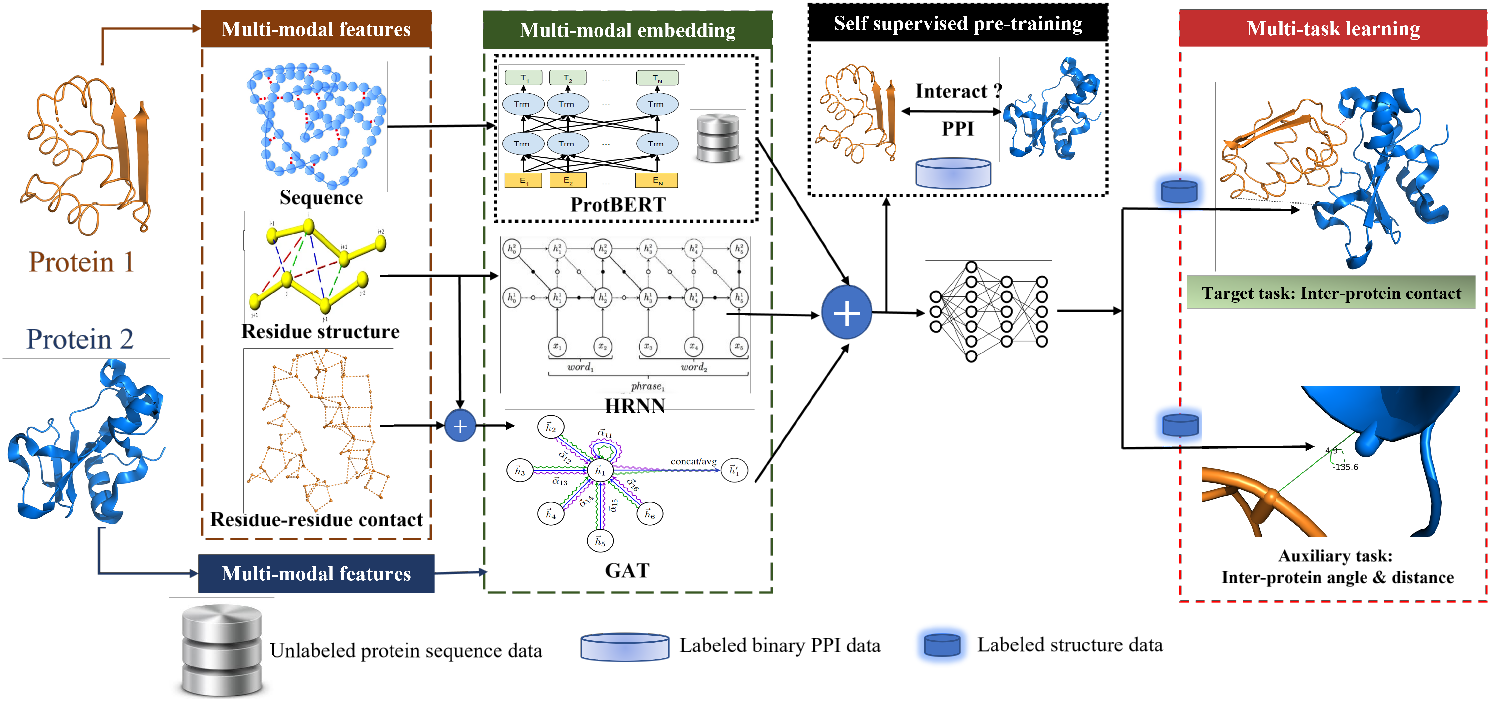
A conceptual overview of using multi-modal representations of proteins to predict contact map, aided by auxiliary tasks. **Left:** Given two proteins, we extract their three modalities as inputs: protein sequences, evolutionary and structural information, with encoders of HRNN, ProtBERT and GAT. **Middle:** Knowledge of protein-protein interaction (PPI) is introduced as an auxiliary task via pre-training. **Right:** 2D geometry prediction (distances and angles) is introduced as an auxiliary task to the target task (inter-protein contact prediction) through multi-task learning / multi-objective optimization.

In the first step to explore auxiliary tasks, we pre-train multi-modal representations to predict protein-protein interactions (PPIs). For model architectures, we replace the contact predictor with a global pooling function and then classifier. PPI models have been trained for 300 epochs with all aforementioned protein modalities as inputs, and with the binary cross entropy loss function, as the warm start for inter-contact prediction since these two tasks do not share datasets.

In the next auxiliary task of distance and angle (2D geometries) prediction, we replace the contact predictor with distance/angle predictors of the similar architecture. We introduce this auxiliary task via multi-task learning (without exploring the weights across the cross entropies across tasks).

### Assessment

We evaluate the performance with the area under the precision-recall curve (AUPRC) since in practice contacts are minorities among non-contacts [9]). To further understand what data samples benefit more from multi-modalities, we split the test protein assemblies based on protein docking difficulty levels [14] quantified by iRMSD: the interface root mean squared difference between unbound and bound protein complex structures. We classify them into three levels of rigid, medium difficult/flexible, and difficult/flexible cases.

### Comparison

The two methods we compare to are ClusPro [22] an FFT-based rigid-body docking method and [13] a GCN neural network-based machine learning method. For ClusPro, the indicators in the residue pair-wise distance matrix used to calculate the AUPRC is the smallest distance between any two heavy atoms across the two residues. The AUPRC of a random classifier for the target task of contact prediction is 9.1% for training and validation dataset (due to under-sampling of negative residue pairs) and 0.1% for the test set.

## 3 Results

### 3.1 Multi-Modal Protein Representations

Figure 2 shows the average validation and test AUPRC scores (%) (with normalized scores via divided by random performance shown in Appendix C. Notice that the scope of validation AUPRC differs a lot from the test; maximum AUPRC for validation and test are 47.6% and 2.15% respectively, due to the different positive/negative ratio of labels (please also refer to the random baselines in red). We observe that, by progressively adding modalities, from protein sequences with HRNN (**H**), then evolutionary profiles with a BERT-like protein language model (**HB**), and finally structure-aware intra-contact maps with GAT (**HBG**), we improve the validation AUPRC accordingly as well as the test. With respect to model **H**, **HBG** improves AUPRC by 7.69% for the validation and 18.75% for the test set. Docking approach (top-1 prediction) compared to machine learning had worse overall performances but relatively high average due to few outliers. We note that the test set is not necessarily when developing the docking method (including its scoring functions).

**Figure 2:**
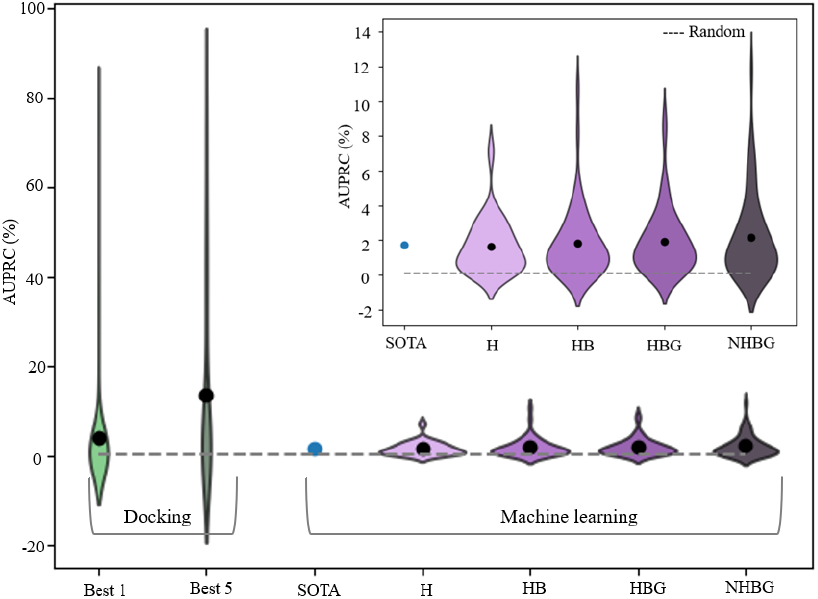
AUPRC (%) distribution of test set. Random score (in grey) shows the positive residual contact base-line for the dataset.

We later also optimize the contact predictor architecture during the study (**NHBG**), mainly by rationally reordering the linear and non-linear layers, which leads to the currently best test performance 2.15%. Our maximum test AUPRC also surpass the SOTA test AUPRC score by 26.47% [13] which only utilizes the single modality of intra-contact maps as inputs. The empirical evidences enable us to conclude that, involving additional protein modalities as inputs benefits interface prediction.

#### More analyses

In Figure. 3, we showed the change of AUPRC for each test protein pair. (1) AUPRC improvement by model **HB** with respect to **H**, due to the addition of a protein language model, is described in Figure 3a. It shows that 83.3% of difficult, 73.3% of mid difficult and 41.4% of rigid PPI samples have improved AUPRC. The protein language model in model **HB** provides global evolutionary information and is less impacted by local sensitivity, which specifically helps the difficult/flexible and medium difficult PPI category. (2) Whereas model **HB** uses multi-views of 1D modality, model **HBG** uses both 1D and 2D modalities. Figure 3b shows that 66.6% of difficult, 33.3% of medium difficult and 69% of rigid PPI samples show AUPRC improvement when using both residue and structure information. Though global evolutionary information shows a significant improvement, the structural data helps the rigid cases the most. Although it loses some of the information about the 3D complex structure, the 2D intra-contact maps provide robust representations by showing less sensitivity to conformational differences between predicted and actual structures.

**Figure 3:**
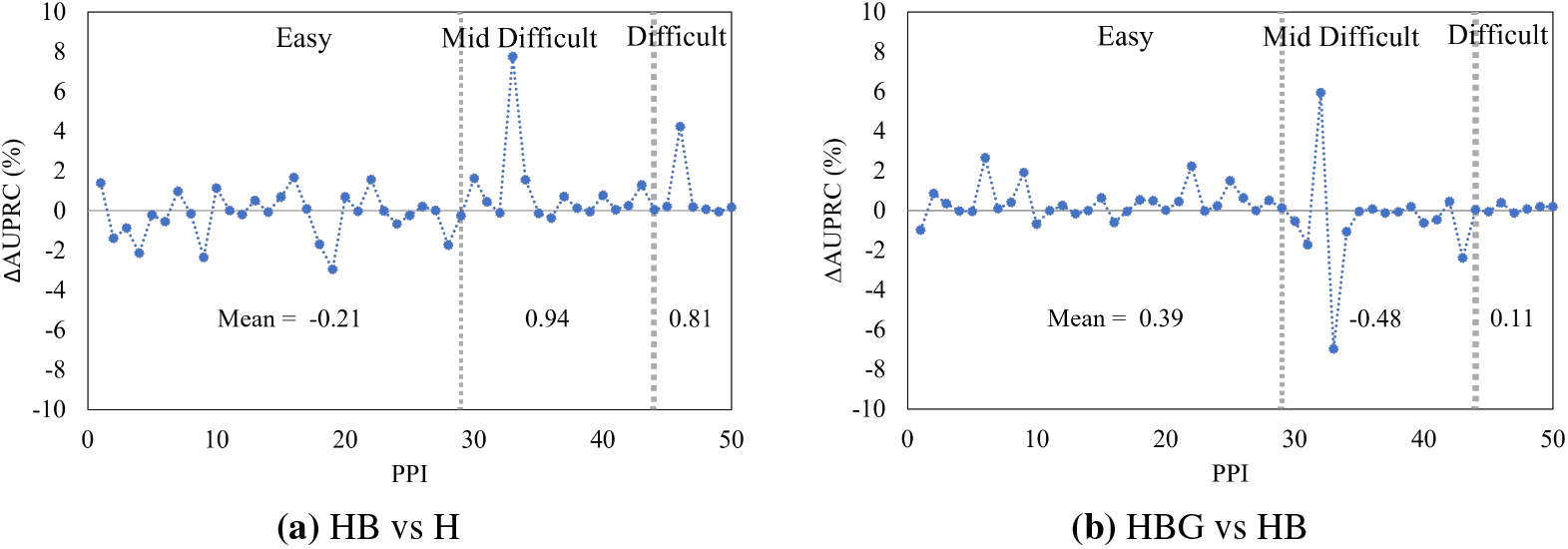
AUPRC improvement with additional modalities on increasingly difficult test PPI levels. We divide the PPI into 3 groups: easy, mid-difficult and difficult group

### 3.2 Auxiliary Task Learning via Pre-Training or Multi-Tasking

The validation and test AUPRC scores average from evaluated protein complexes, are shown in Figure 7. Clearly, we do not observe the performance gains from any auxiliary tasks. There are several conjectures for the negative results. First, simultaneously predicting more fine-grained contact distances and angles presumably is informative for contact prediction but it is not. It might result from the limited interface data: enforcing modals to output more complicated (pseudo) predictions while with the same restricted amount of data, might even hurt generalizability.

**Figure 4:**
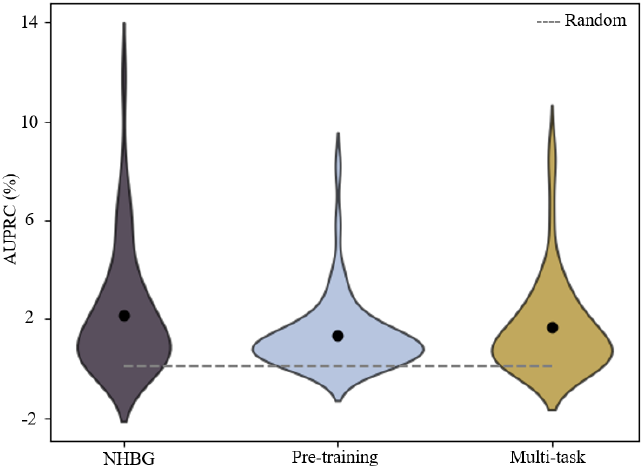
Average AUPRC % of test set over auxiliary task optimization models

**Figure 5:**
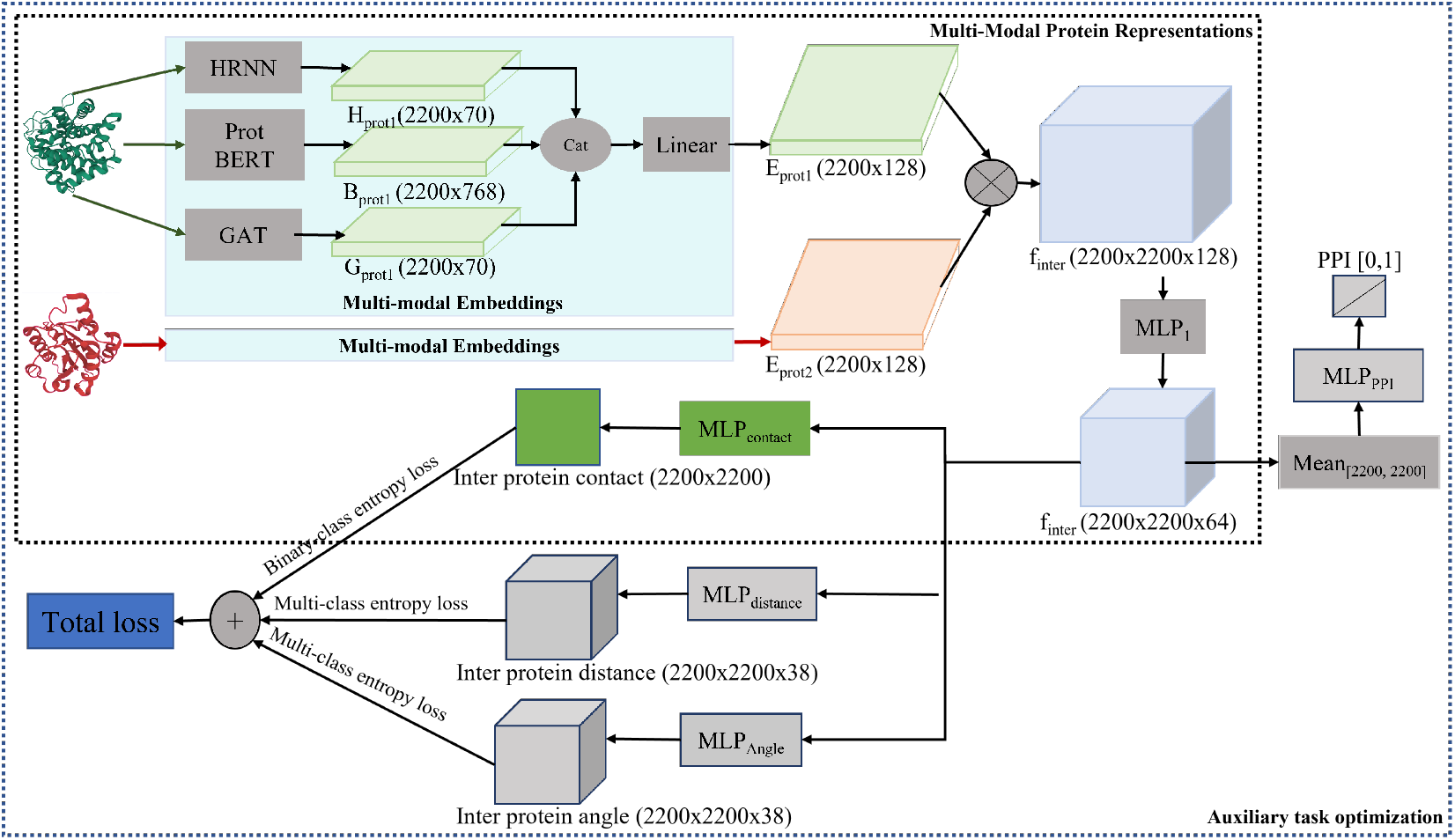
A detailed framework of our model architecture using separate encoders, a task-shared common MLP layer, and task-specific MLP layers tuned by cumulative loss of all tasks.

**Figure 6:**
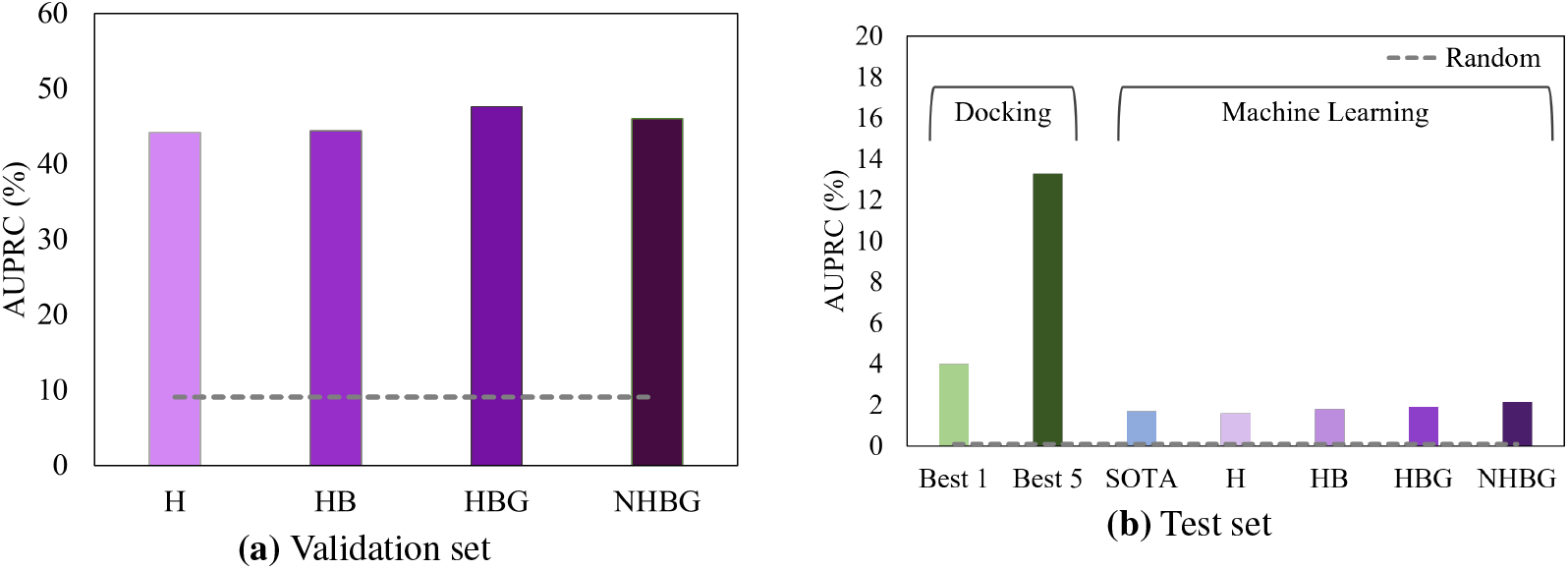
Average AUPRC (%) of validation and test set. Random score (in grey) shows the positive residual contact baseline for each dataset.

**Figure 7:**
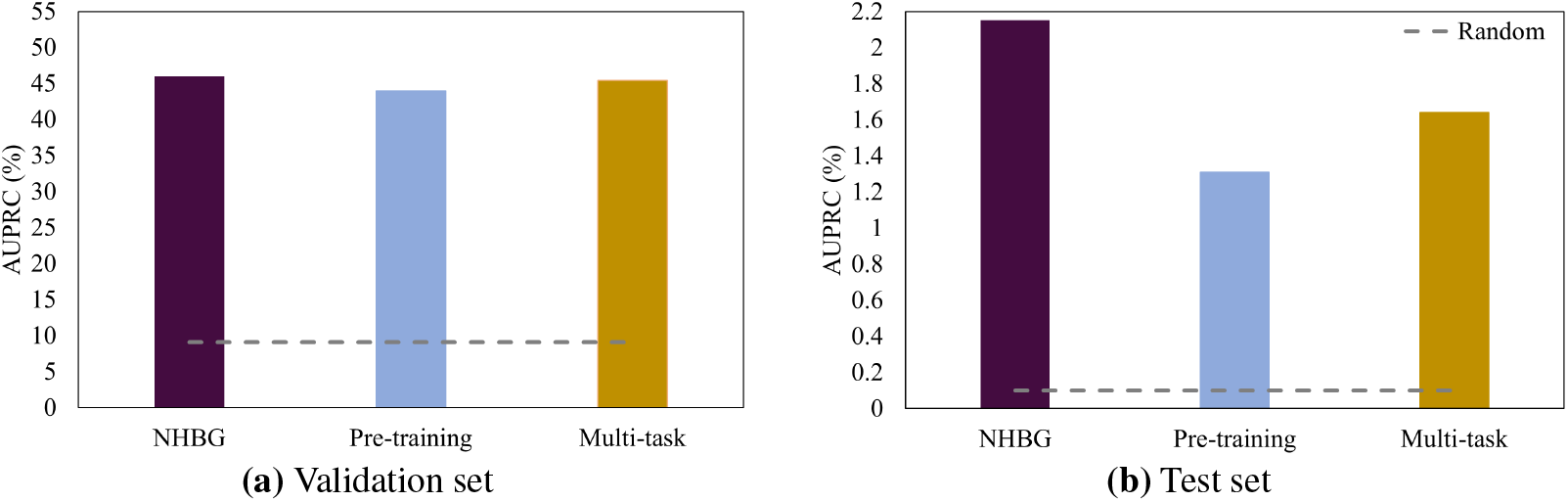
Average AUPRC (%) of validation and test set over auxiliary task optimization models.

Second, PPI as the consequence of inter-protein contacts also does not benefit. Although PPI data are more accessible, the information encoded in the interaction (binary outcome, interacting or not) is overly simplified to indicate inter-contact maps (with the complexity proportional to multiplication of protein pairs’ lengths). This indicates both data quantity and quality are prerequisite for constructing synergistic auxiliary tasks, which is nontrivial for complicated protein complex data.

Lastly, analyses similar to Section 3.1 is conducted with no significant performance difference on data across all difficulty categories. Results can be found in Appendix C.

## 4 Conclusions

In this paper we study how the multi-modal protein data and auxiliary predictive tasks can impact inter-protein contact prediction. We first progressively add different modalities as sequence, evolutionary and structural information. Our results show that multi-modalities provide significant benefits over single modality to predict inter-contacts. We next introduce auxiliary tasks of protein-protein interaction prediction (via pre-training) and inter-contact distance and angle prediction (via multi-task learning). We numerically find that neither improves predictions, indicating that constructing and incorporating synergistic auxiliary tasks for predicting protein interfaces remains challenging.

## Acknowledgments and Disclosure of Funding

We thank the National Science Foundation for funding the study (CCF-1943008) and Texas A&M High Performance Research Computing for providing computational support.

## Appendix

## A Details on Data Curation

Extra efforts are paid to sanitize the DB5 unbound–bound structure data. Specifically if a protein has multiple chains, in [13] residues in unbound receptor/ligand proteins were indexed according to some undisclosed order of chain concatenation. The chain concatenation order is determined by picking the one with the best scores of Needleman–Wunsch alignment between the [13] sequences converted from hydrophobicity [23] vertex features (residues with hydrophobicity index −3.5 are viewed as glutamic acids) and the DB5 sequences among all permutations of chains. In the meantime, the alignment also indicated the correspondence of residues between the [13] sequences and the DB5 cleaned PDB file sequences. To generate the alignment between residues in the bound and unbound sequences, for receptor/ligand respectively, we also used Needleman-Wunsch alignment between the sequences from the unbound and bound structure files.

For labeling in the target task, we retrieve the ground-truth contact labels *C*_label_ ∈ {0, 1}^*L*_1_×*L*_2_^ from the bound structures and assign them nonzero values of ones only if the shortest distances between non-hydrogen (heavy) atoms across two residues are below the threshold of 6Å [13, 24, 25, 26]. For labeling the auxiliary task, we retrieved the 2D geometries from bound structures and discretize them. Distance bins are (0-2Å, 2-2.4Å, 2.4-2.8Å, …, 19.6-20Å, and >20Å) plus two additional bins: no contact (distance >20Å) and false mask for missing atom/residue in the diagonal line of contact map. All the angles have been divided into 36 bins ranging from −*π* to *π* plus no contact and false mask.

## B Details on Model Architectures

## C Additional Results

**Table 1:** Normalized AUPRC divided by random performance.

| Model | Validation | Test |
| --- | --- | --- |
| H | 4.85 | 16 |
| SOTA | - | 17 |
| HB | 4.88 | 18 |
| HBG | 5.23 | 19 |
| NHBG | 5.05 | 21.5 |

**Figure 8:**
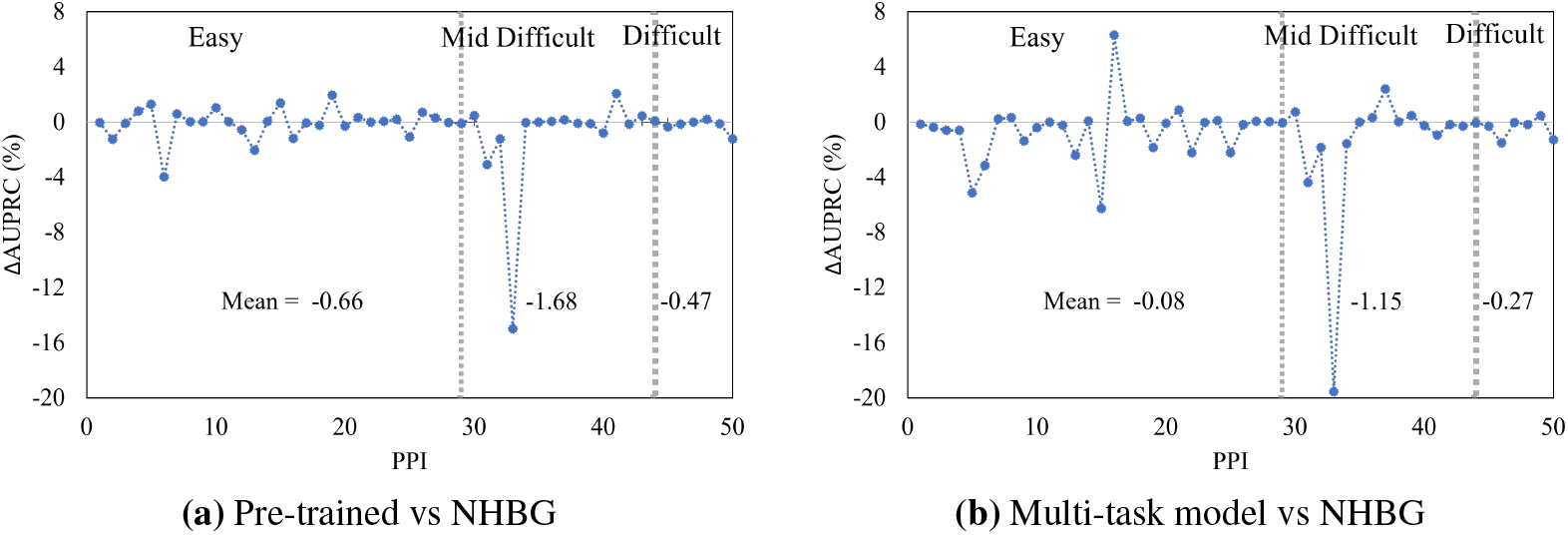
AUPRC improvement with auxiliary task optimization compared to new HBG model. We divide the test PPI into 3 groups based on increasing docking difficulty / conformational changes.

